# Spatial Conformation of Pancreatic Cancer Is Associated with Disease Recurrence after Curative-Intent Total Neoadjuvant Therapy

**DOI:** 10.1101/2025.11.03.686383

**Authors:** Luis H. Cisneros, Merih D. Toruner, Zafar Siddiqui, Andrea V. Maraone, Miranda Lin, Alex Xiao, Cornelius Thiels, Mark J. Truty, Ben George, Khalid Jazieh, Rob McWilliams, Carlo C. Maley, Chris Hartley, Rofyda Elhalaby, Paul Dizona, Qian Shi, Martin E. Fernandez-Zapico, Ryan M. Carr

## Abstract

Pancreatic ductal adenocarcinoma (PDAC) frequently recurs after total neoadjuvant therapy (TNT) and curative-intent resection. Traditional histopathologic response assessments demonstrate that major pathologic response (<5-10% viable residual cancer) is associated with favorable outcomes. However, most TNT cases achieve only a minor response. We investigated whether the spatial organization of residual PDAC encodes clinically meaningful biology beyond residual tumor burden. In a retrospective cohort of 203 resected PDAC patients, all with minor pathologic response, H&E whole-slide images were segmented into cancer and stroma to subsequently quantify spatial composition (e.g., patch size/density, edge density) and configuration (e.g., patch complexity/compactness, spatial intermixing) to model treatment-resistant tumor architecture. Non-response was associated with a more fragmented interface-rich ecology, higher edge density and diversity, and reduced homotypic aggregation, independent of conventional clinicopathologic features. This demonstrates a link between emergent tissue architecture of treated PDAC and therapeutic resistance. Further, two multivariable, spatial risk models were independently associated with disease-free survival (DFS): (1) cancer mean shape index and stromal shape-index variability (high-risk median DFS 7.23 vs 11.57 months; adjusted HR 1.75, p=0.007) and (2) mean stromal area with edge density (high– vs low-risk adjusted HR 1.94, p=0.006), outperforming traditional treatment response assessments. Quantifying residual cancer–stroma topology thus yields independent, prognostic signals in post-TNT PDAC and motivates prospective, spatially informed adjuvant strategies and mechanistic studies of edge habitats and mixing as therapeutic vulnerabilities.

## Introduction

Despite therapeutic advances, pancreatic ductal adenocarcinoma (PDAC) remains among the most lethal cancers, with high rates of recurrence despite extensive total neoadjuvant therapy (TNT) and curative-intent resection^1^. While TNT has improved treatment completion^2,3^, margin-negative resection (R0) rates^4,5^, nodal clearance^6^, pathologic response^7^, and survival outcomes^4,6,8^, more than half of patients develop disease recurrence within two years^9^.

Pathologic treatment response correlates with outcome at the extremes: a major response is favorable^10–15^, whereas most patients achieve only a minor response with heterogeneous courses thereafter^9^. Within this large majority with residual disease, current assessments emphasize how much tumor persists rather than how it is organized, relying on semi-quantitative or subjective scoring that can vary across observers^12,13^. These limitations constrain risk stratification and do not consider spatially informed hypotheses about therapeutic resistance.

Tumors are structured ecosystems^16^ in which cancer and stroma form patches and interfaces^17^ influencing nutrient flow^18–20^, immune access^21–25^, mechanical constraints^26–28^, and routes for invasion^29,30^. After therapy, selection can re-shape this ecology, potentially enriching spatial configurations that enable disease persistence^31^. Conventional pathologic treatment response assessments, capturing residual disease burden, may be enhanced through quantification of emergent topologies (e.g., fragmentation, interface density, homotypic clustering, or shape irregularity) that may encode clinically relevant biological information related to therapeutic resistance, disease persistence, and risk of relapse.

Here, focusing deliberately on the challenging and clinically prevalent cases of PDAC with minor pathologic responses after TNT and resection, we ask whether residual cancer-stroma spatial organization predicts disease-free survival (DFS) beyond standard clinicopathologic variables and gross disease extent. Using H&E whole-slide overlays to delineate cancer and stroma, we operationalize a computational pipeline^32^ to quantify composition (e.g., patch size/density, edge density) and configuration (e.g., patch complexity/compactness, spatial intermixing) at patch and landscape scales, and we develop multivariable spatial risk models. By reframing treated PDAC as a post-therapy landscape, rather than only focusing on disease burden, we aim to provide a quantitative, observer-independent description of the microecology that persists after TNT, furnish prognostic insights in the context of most cases with minor pathologic responses, and generate mechanistic hypotheses about interface habitats and mixing as potential therapeutic vulnerabilities.

## Results

### Study Population, Surgical, and Pathological Outcomes

The study cohort consists of 297 samples for 203 patients. Each patient had one or two WSIs analyzed. Of these, 124 had PDAC recurrence, 60 of which had two WSIs included in the analysis. Eighty did not have recurrence within up to 150 months follow up time, 34 of which had two WSIs analyzed.

All 203 patients received TNT. Among patients who undergo TNT, a minority (approximately 20%^9^) achieve a major pathologic response by College of American Pathologists (CAP) criteria^33^, which carries a well-established favorable prognosis^10–15^. Conversely, all patients in the present cohort achieved a minor pathologic response with CAP grade 2 or 3 indicative of partial response and no response, respectively. The mean follow-up time was 18.9 months. Demographic and clinicopathologic characteristics are summarized in **Table 1**. The mean age at surgery was 65.4 years with a relatively balanced distribution of sex (47.3% female). Most patients had clinical stage IB (30.5%), IIB (25.5%), or III (26.0%) disease at diagnosis.

**Table 1.** Demographics and Clinicopathologic Factors.

|  | Overall (N=203) |
| --- | --- |
| **Age at Surgery** |  |
| □□□Mean (SD) | 65.4 (9.2) |
| □□□Median (Range) | 66.5 (39.4, 85.3) |
| **Race** |  |
| □□□White | 192 (95.5%) |
| □□□Non-White | 9 (4.5%) |
| **Gender** |  |
| □□□Female | 96 (47.3%) |
| □□□Male | 107 (52.7%) |
| **Clinical Stage at Diagnosis** |  |
| □□□IA | 23 (11.5%) |
| □□□IB | 61 (30.5%) |
| □□□IIA | 12 (6.0%) |
| □□□IIB | 51 (25.5%) |
| □□□III | 52 (26.0%) |
| **Type of Surgical Resection** |  |
| □□□Distal Pancreatectomy | 45 (22.2%) |
| □□□Pancreaticoduodenectomy | 126 (62.1%) |
| □□□Total Pancreatectomy | 32 (15.8%) |
| **Time from Diagnosis to Surgery (months)** |  |
| □□□Mean (SD) | 8.6 (3.2) |
| □□□Median (Range) | 8.2 (1.8, 19.4) |
| **Histological Grade** |  |
| □□□1 - Well-differentiated | 4 (2.0%) |
| □□□2 - Moderately differentiated | 138 (68.0%) |
| □□□3 - Poorly differentiated | 57 (28.1%) |
| □□□4 - Undifferentiated | 2 (1.0%) |
| □□□X - Cannot be determined | 2 (1.0%) |
| **Local Invasion** |  |
| □□□Confined to Pancreas | 56 (28.4%) |
| □□□Duodenal Wall | 23 (11.7%) |
| □□□Peripancreatic Soft Tissues | 64 (32.5%) |
| □□□Multiple | 47 (23.9%) |
| □□□Other | 7 (3.6%) |
| **Pathologic Response (CAP)** |  |
| □□□2 - Partial Response | 156 (82.2%) |
| □□□3 - No Response | 34 (17.9%) |
| **Resection Outcome** |  |
| □□□R0 | 174 (86.1%) |
| □□□R1 | 27 (13.4%) |
| □□□R2 | 1 (0.5%) |
| **Lymphovascular Invasion** |  |
| □□□Yes | 27 (14.4%) |
| □□□No | 159 (84.6%) |
| **Perineural Invasion** |  |
| □□□Yes | 110 (57.3%) |
| □□□No | 80 (41.7%) |
| **Pathologic T Stage** |  |
| □□□1a | 7 (3.5%) |
| □□□1b | 4 (2.0%) |
| □□□1c | 35 (17.3%) |
| □□□2 | 102 (50.5%) |
| □□□3 | 44 (21.8%) |
| □□□4 | 10 (5.0%) |
| **Number of Nodes Removed, Median (Range)** | 20.0 (1.0, 71.0) |
| **Number of Positive Nodes, Median (Range)** | 0.0 (0.0, 17.0) |
| **Pathological N Stage** |  |
| □□□N0 (0 LN+) | 140 (69.0%) |
| □□□N1 (1-3 LN+) | 45 (22.2%) |
| □□□N2 (4+ LN+) | 18 (8.9%) |
| **Neoadjuvant Treatment** |  |
| □□□Chemo and RT with radiosensitizing chemo | 149 (73.4%) |
| □□□Chemo and RT w/o radiosensitizing chemo | 2 (1.0%) |
| □□□Chemo only | 43 (21.2%) |
| □□□RT with radiosensitizing chemo | 9 (4.4%) |

Most subjects underwent pancreaticoduodenectomy (62.1%) and achieved an R0 resection (86.1%). The median time from diagnosis to surgery was 8.2 months (range 1.8-19.4 months). Histological grading indicated that 68.0% were moderately differentiated, while 28.1% were poorly differentiated.

Pathologic staging by AJCC v.8 criteria showed that 50.5% were classified as T2 tumors, while 21.8% and 5.0% were T3 and T4, respectively. Nodal involvement was observed at 31.0%, with 22.2% classified as N1 and 8.9% as N2. Perineural invasion (PNI) and lymphovascular invasion (LVI) were identified in 57.3% and 14.4% of cases, respectively. These findings underscore the heterogeneity of tumor characteristics and underline the persistent burden of disease in this cohort despite TNT and surgical intervention.

### Clinicopathologic Features and Disease-Free Survival

Univariate analysis identified several clinicopathological factors associated with DFS (**Supplementary Table 1**). Pathologic T (ypT) stage (p<0.001) (**Figure 1A**), pathologic N (ypN) stage (p=0.002) (**Figure 1B**), and LVI (p=0.048) were significantly associated with DFS. Other factors, such as age, gender, histologic grade, and time from diagnosis to surgery, were not significantly associated with time to recurrence.

**Figure 1.**
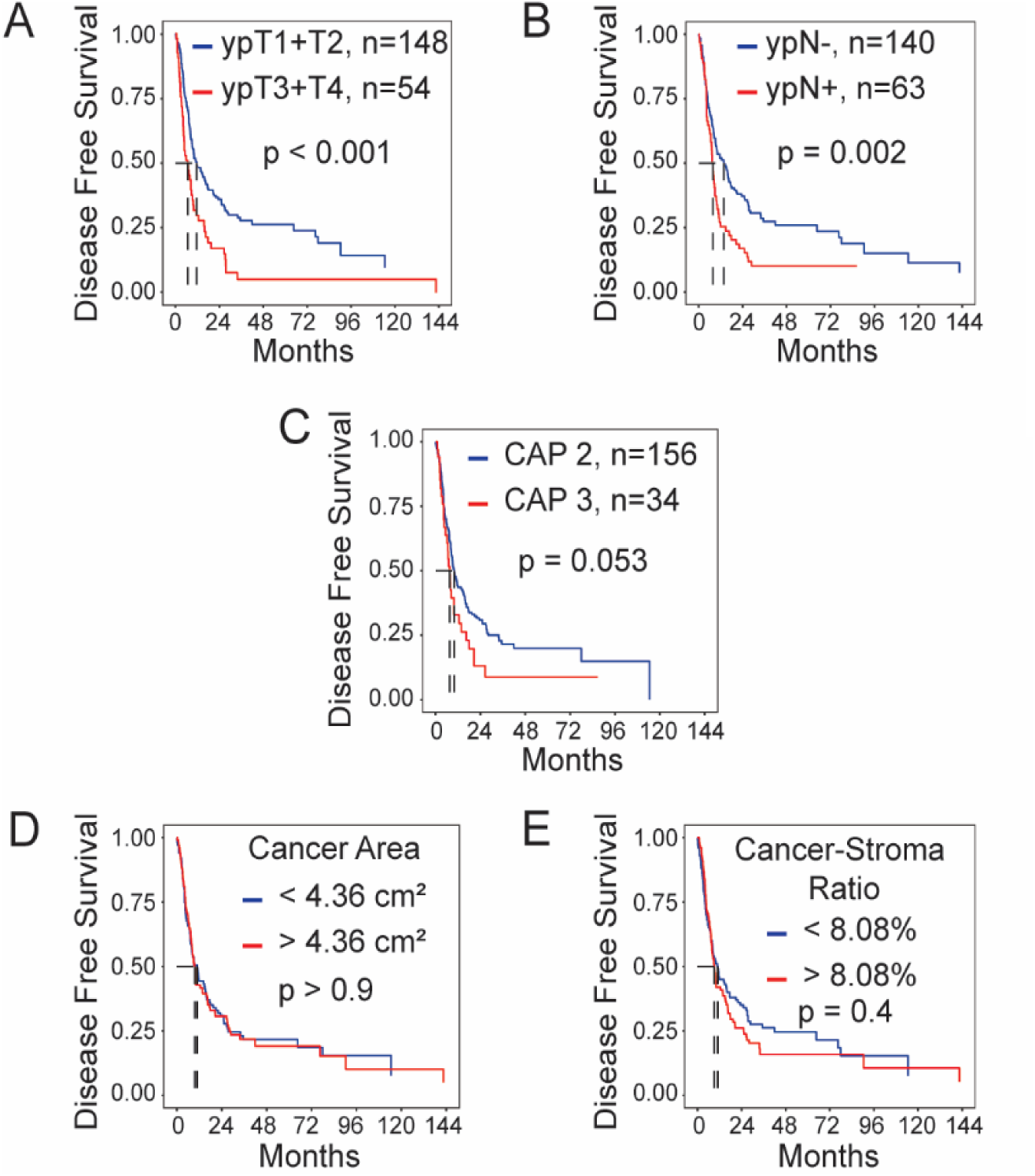
Clinicopathologic features associated with disease-free survival (DFS). Kaplan-meier curves stratifying DFS by (A) pathologic tumor (ypT) stage, (B) pathologic nodal (ypN) stage, (C) College of American Pathologists (CAP) tumor regression grade 2 (partial response) versus grade 3 (no response), (D) area of residual cancer within the tumor bed, and (E) ratio of cancer area to stroma area within the tumor bed.

Outcomes between cases with CAP grades 2 and 3 were not significantly different (**Figure 1C**). Given the subjective nature of CAP assessments by pathologists and risk of inter-observer variability^12,13^, we assessed whether quantification of residual cancer could stratify outcomes in this cohort. Interestingly, quantifying the area occupied by cancer within the residual tumor was not associated with DFS (p=0.95) (**Figure 1D**). The area occupied by cancer relative to the stroma area was also quantified much like published treatment response scores.^13^ This ratio was similarly not associated with outcomes (p=0.40) (**Figure 1E**). While quantifying the amount of the residual tumor failed to stratify patient outcomes, this led us to explore whether the spatial composition and configuration might offer more significant biological and clinical insights into residual disease.

### Spatial Pathological Cohort Characterization

Using PathExplore™ PDAC tissue overlays reduced to cancer and stroma patches, we summarized tissue composition (abundance and density of patches) and configuration (their spatial arrangement) at the cohort level and related these distributions to CAP treatment response categories (**Figures 2 and 3**). At baseline, stromal patches were consistently larger and more dispersed than cancer patches (**Figure 2A-B**). When stratified by response, mean cancer patch area trended lower in CAP2 than CAP3, while variability in patch area separated groups for both stroma and cancer, with higher variability in CAP3 (**Figure 2A-B**). Extremes of patch extent, captured by minimum/maximum Feret diameter^34^, did not differ by response (**Figure 2C–D**). Patch density differed by response for cancer but not for stroma, consistent with more numerous, smaller cancer islands in CAP3 (**Figure 2E**). Edge density, the total interface length per area, was markedly higher in CAP3, indicating a more fragmented, interface-rich landscape when tumors failed to respond (**Figure 2F**). Finally, diversity indices (Shannon and Simpson diversity indices) were elevated in CAP3, reinforcing that non-responders harbor more compositionally and spatially diverse landscapes (**Figure 2G-H**).

**Figure 2.**
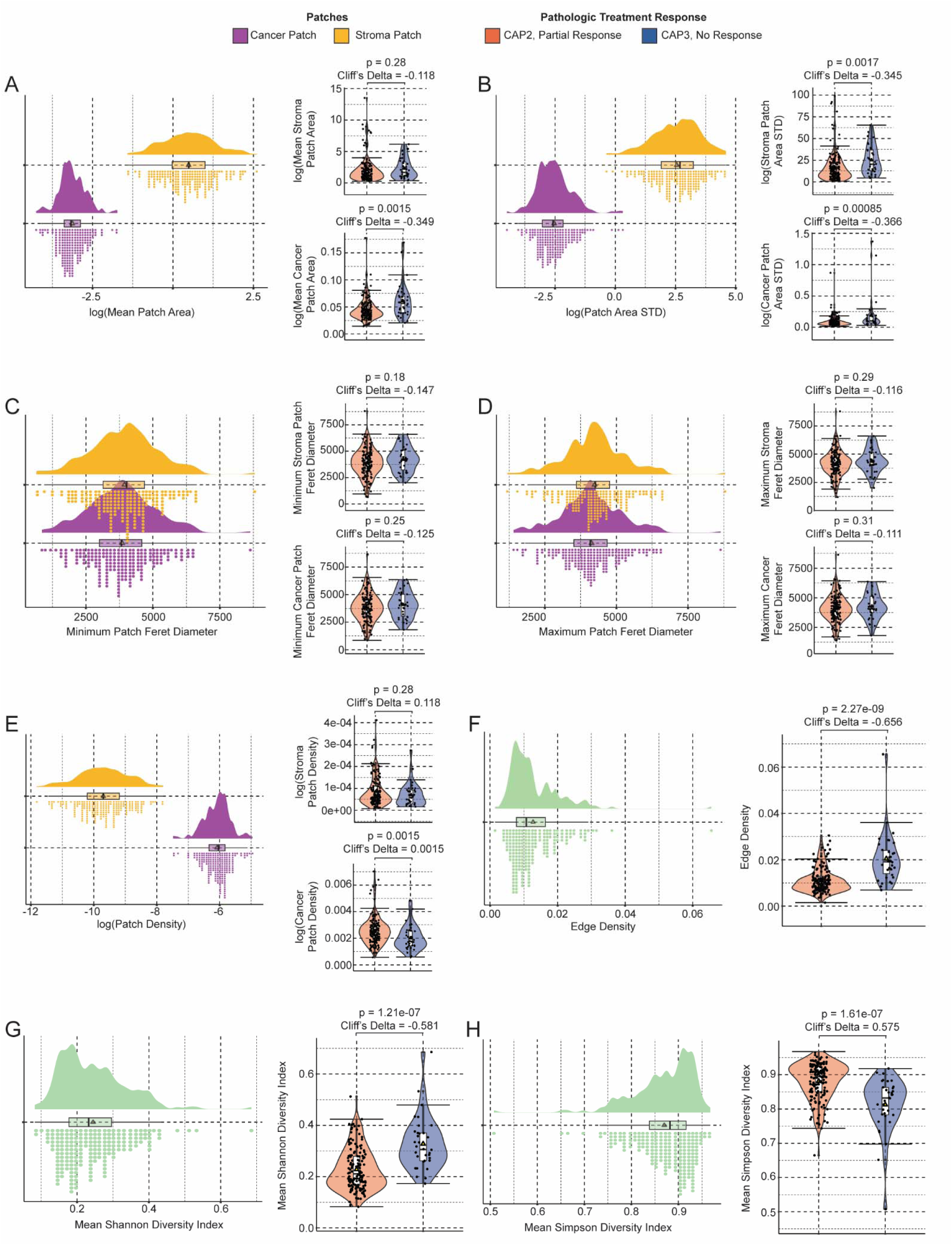
Cohort-level composition and divesity spatial pathology feature sumaries and response-stratified comparisons. Cohort-level distributions and CAP response-stratified comparisons for spatial features. For panels that are patch-level and compartement-specific (A-E), the left plots show cohort distributions by tissue compartment, cancer patches (purple) and stroma patches (yellow), with kernel density, boxplot, and rug marks. The right plots show per-patient values summarized by CAP pathologic treatment response using violin plots with overlaid points. CAP2 (partial response) is in red and CAP3 (no response) is in blue. Panel (F) shows landscape edge density (whole-slide composition metric). Panels (G) and (H) show mean Shannon diversity index and mean Simpson diversity index, respectively. STD = standard deviation

**Figure 3.**
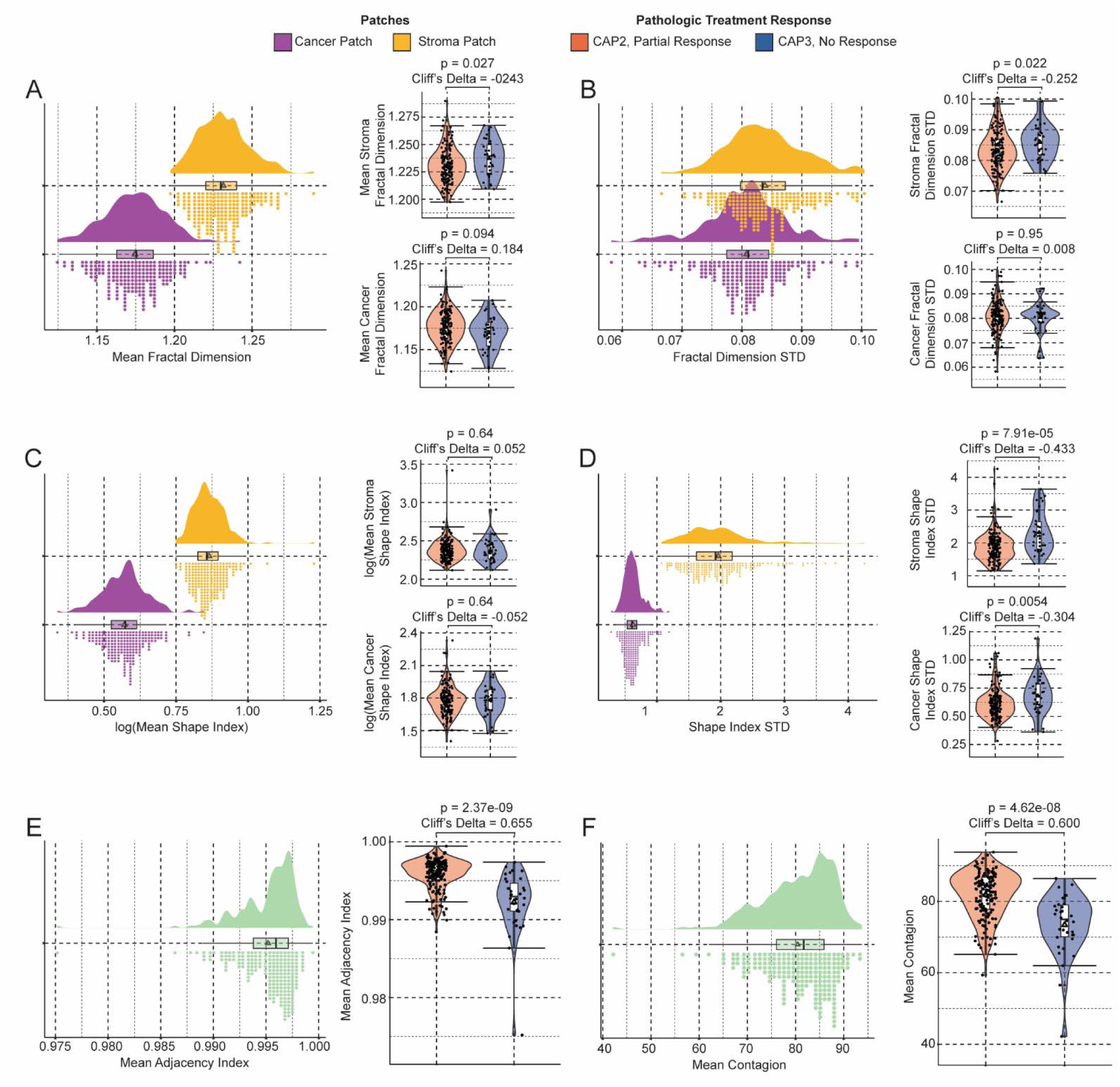
Cohort-level configuration spatial pathology feature summaries and response-stratified comparisons. Panels A-D summarize patch-level, compartment-specific geometry for cancer (purple) and stroma (yellow). For eatch metric, the left plot shows the cohort distribution by compartment using kernel density, boxplot, and rug marks. The right plot shows per-patient summaries stratified by CAP pathologic treatment response with violin plots and overlaid points. CAP2 (partial response) is in red and CAP3 (no response) is in blue. Panels E-F display whole-slide configuratoin metrics with the same layout. € mean adjacency index (homotypic contacts) and (F) mean contagion (landscape aggregation).

Metrics of configuration also exhibited separation by treatment response (**Figure 3**). Fractal dimension, a boundary complexity measure, showed higher mean stromal patch complexity and higher stromal variability in CAP3, while cancer fractal metrics were not discriminatory (**Figure 3A-B**). However, there was some association between decreased fractal dimension and poorly differentiated disease (**Supplementary Figure 1**) Mean shape index (compactness) did not differ by response (**Figure 3C**), but shape index variability did. Larger shape index variability was seen in CAP3 stroma and cancer (**Figure 3D**). Adjacency index (like-with-like contact) and contagion (clumping) were both lower in CAP3, pointing to reduced homotypic aggregation and greater intermixing of cancer and stroma when treatment was ineffective (**Figure 3E-F**). None of these spatial features were consistently associated with common clinicopathologic features of disease such as differentation or pathologic T or N staging (**Supplementary Figures 1-3**).

In sum, across scales, non-response was characterized by (1) higher variability in patch geometry (area, fractal dimension, shape index), most prominently within stroma, and (2) greater fragmentation and mixing (higher edge density, lower adjacency/contagion, higher diversity).

Figure 4 presents a cohort-level heatmap of spatial pathology features across individual patients. Two patient groups emerge when maximizing the average silhouette width for k-means^35^ using all the features (**Supplementary Figure 4**). Cluster 1 exhibits a fragmented, interface-rich phenotype characterized by a higher landscape edge density and diversity indices, lower contagion and homotypic adjacency, smaller mean patch sizes with greater patch density and modestly higher variability in shape/fractal metrics. Cluster 2 shows the reciprocal pattern: larger mean patch sizes with fewer patches, and overall, a coarser, more aggregated tissue organization. The corresponding clinical features are shown color bars on top of the heatmap. Of these, CAP3 samples are significantly enriched in Cluster 1 (p-value < 0.0004), per a standard Chi-square goodness of fit test, while CAP2 are marginally enriched in Cluster 2 (p-value = 0.017). While Cluster 1 and Cluster 2 cases did not have statistically significant different DFS, these cohort-level patterns support the central premise that emergent spatial organization of the residual cancer–stroma ecosystem captures treatment resistance beyond gross burden alone. This motivated subsequent statistical modeling investigating the clinical significance of these spatial measures.

**Figure 4.**
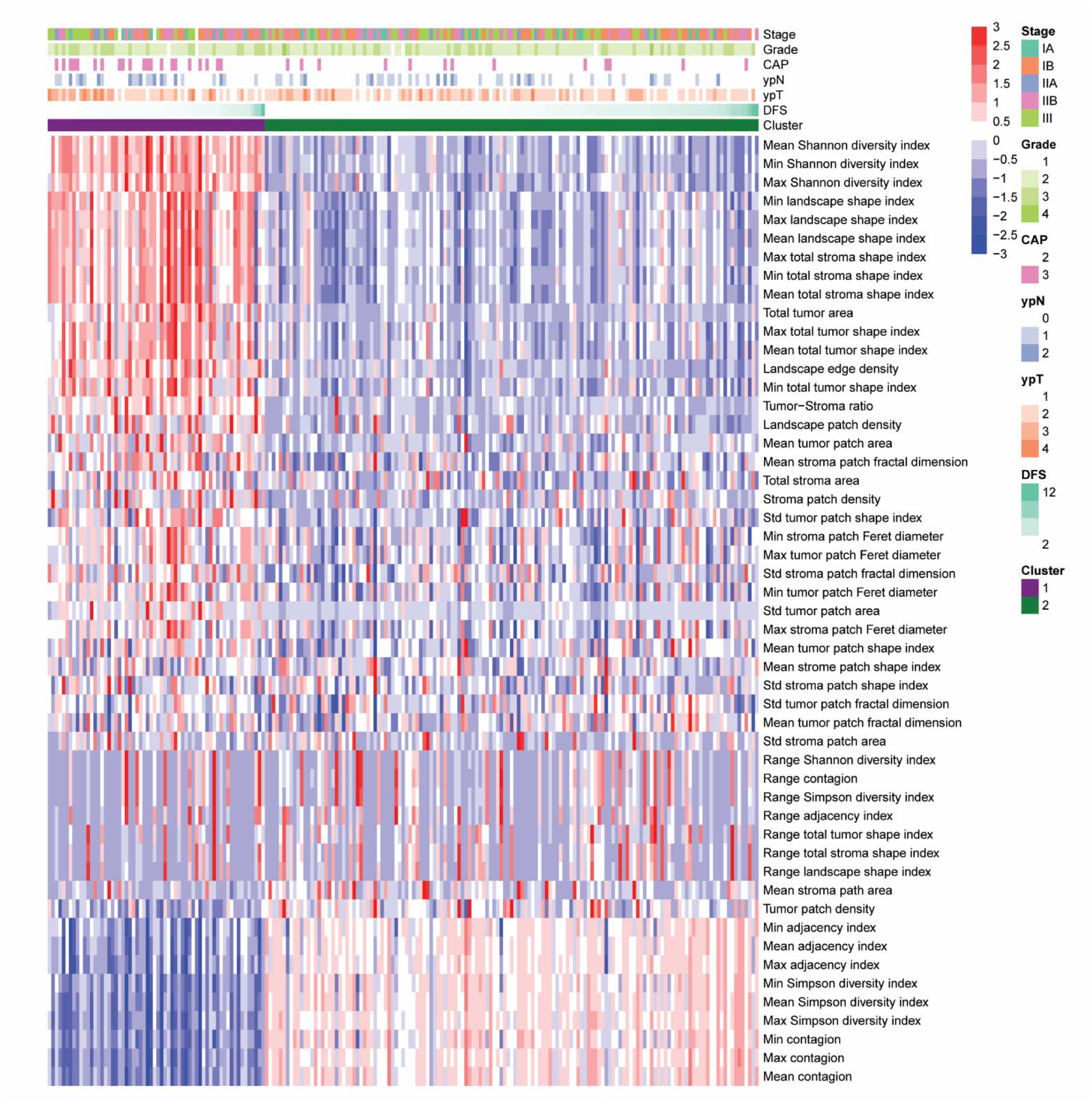
A cohort-level heatmap of z-scored spatial pathology features (rows) across individual patients (columns) with unsupervised hierarchical clustering of patients (k-means). The color scale reflects relative deviation from the cohort mean. CAP = College of American Pathologist pathologic treatment response; ypN = pathologic lymph node status; ypT = pathologic tumor status; DFS = disease free survival.

### Spatial Pathological Feature Selection

Cross-correlation between all 31 variables reveals a significant overlap for 15 variables, which group into 5 different clusters. Each cluster is comprised of 2 to 6 variables for which the confidence interval of the Pearson’s cross-correlation overlaps the value 1.0, indicating that full correlation cannot be statistically discarded (and thus they are likely to carry the same information) (**Supplementary Figure 5**). For each of the clusters, the variable that has the lowest p-value in a Wilcox test that distinguishes the recurrence status was chosen as the best representative, reducing the total number of variables to 20.

Spatial variables were then evaluated to identify those most strongly associated with DFS. Survival analyses identified five features with p-values <0.20: natural log (ln) of the standard deviation (SD) of the stroma patch area, ln(edge density), mean cancer shape index (CSI), ln(SD stroma shape index [SSI]), and the ln(minimum CSI) (**Table 2**). To further evaluate the potential biological and clinical significance of these features, statistical models were developed.

**Table 2.** Univariate Association Between Spatial Pathology Features and DFS (Cox Regression and Spline Models)

| Spatial Variables | Cox Model |  |  |  | Spline Analysis |  |  |
| --- | --- | --- | --- | --- | --- | --- | --- |
|  | HR | Conf.low | Conf.high | p-value | Overall p-value | Linear p-value | Non-linear p-value |
| Ln(cancer area mean) | 0.93 | 0.62 | 1.38 | 0.7089 | 0.4375 | 0.8057 | 0.3813 |
| Ln(cancer area sd) | 1.03 | 0.81 | 1.30 | 0.8069 | 0.7174 | 0.8396 | 0.6212 |
| Ln(stroma area mean) | 1.27 | 1.03 | 1.56 | **0.0224** | 0.0385 | 0.0200 | 0.3638 |
| Ln(stroma area sd) | 1.16 | 0.98 | 1.37 | 0.0929 | 0.1866 | 0.1231 | 0.5794 |
| patch_density_1e4 | 0.99 | 0.87 | 1.13 | 0.8807 | 0.5252 | 0.9109 | 0.5822 |
| patch_density_1_1e3 | 1.05 | 0.90 | 1.23 | 0.4986 | 0.3383 | 0.4407 | 0.3874 |
| Ln(stroma patch density) | 0.79 | 0.64 | 0.97 | **0.0224** | 0.0385 | 0.0197 | 0.3681 |
| Ln(edge density) | 1.40 | 1.04 | 1.87 | **0.0263** | 0.0636 | 0.0382 | 0.4208 |
| Cancer shape index mean | 0.27 | 0.08 | 0.95 | **0.0407** | 0.1301 | 0.0383 | 0.6248 |
| Cancer shape index sd | 0.81 | 0.25 | 2.62 | 0.7275 | 0.5199 | 0.7266 | 0.5609 |
| Ln(stroma shape index, mean) | 3.88 | 0.19 | 77.46 | 0.3754 | 0.1266 | 0.3265 | 0.1560 |
| Ln(stroma shape index sd) | 2.38 | 1.26 | 4.49 | **0.0075** | 0.0383 | 0.0078 | 0.5265 |
| Cancer shape index range | 0.99 | 0.97 | 1.02 | 0.5314 | 0.3964 | 0.6955 | 0.4062 |
| Shannon diversity index mean | 2.63 | 0.51 | 13.53 | 0.2477 | 0.5787 | 0.2729 | 0.7737 |
| Shannon diversity index range | 0.16 | 0.00 | 9.70 | 0.3786 | 0.7299 | 0.4599 | 0.7710 |
| feret_diameter_1_min_1eneg3 | 1.00 | 0.88 | 1.13 | 0.9733 | 0.7942 | 0.9910 | 0.7745 |
| feret_diameter_1_max_1eneg3 | 0.99 | 0.86 | 1.14 | 0.8776 | 0.7579 | 0.8967 | 0.7358 |
| Ln(cancer shape index minimum) | 1.40 | 1.01 | 1.93 | **0.0433** | 0.1008 | 0.0682 | 0.4157 |
| Shape index maximum | 1.00 | 0.99 | 1.02 | 0.7825 | 0.7233 | 0.7862 | 0.7101 |
| Shape index range | 0.99 | 0.96 | 1.02 | 0.5130 | 0.4590 | 0.6471 | 0.5485 |

### Variability in Stromal Shape Index and Mean Cancer Shape Index

Using recursive partitioning and tree regression (RPTR) to stratify patients into high– and low-risk groups, CSI and SSI were identified as significantly discriminatory (**Supplementary Table 2**). Shape index is a metric used to measure the complexity of the shape of a habitat patch, providing insight into how irregular or convoluted the boundary of the patch is compared to a standard geometric shape, such as a square or circle. Shape index is calculated as follows:

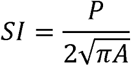

Where *P* represents perimeter of the patch, *A* is the area of the patch. A shape index of 1 is indicative of a perfectly circular patch, representing minimum complexity. Alternatively, the greater that shape index, the more irregular or elongated the patch.

Risk classification was specifically defined by ln(mean CSI) (Figure 5A and B) and ln(SD SSI) (Figure 5C and D). High-risk disease was defined as ln(mean CSI) ≤ 1.775 and ln(SD SSI) > 0.651. High-risk patients (n=63) had a median DFS of 7.23 months (95% CI: 5.19-9.63) compared to 11.57 months (95% CI: 9.24-17.92) for low-risk patients (HR=1.82; 95% CI: 1.31-2.52; p=0.0003) (Figure 5E). After adjusting for the ypT and ypN stages, lymphovascular invasion/perineural invasion, resection outcome, and local invasion by multivariable Cox regression analysis (**Supplementary Table 3**), spatial indices remained independently prognostic for DFS. High-risk classification was associated with a 1.75-fold increased risk of recurrence (HR=1.75, 95% CI: 1.17-2.63, p=0.007). Sensitivity analysis also included other clinicopathologic covariates, which resulted in consistent results (**Supplemental Table 4**).

**Figure 5.**
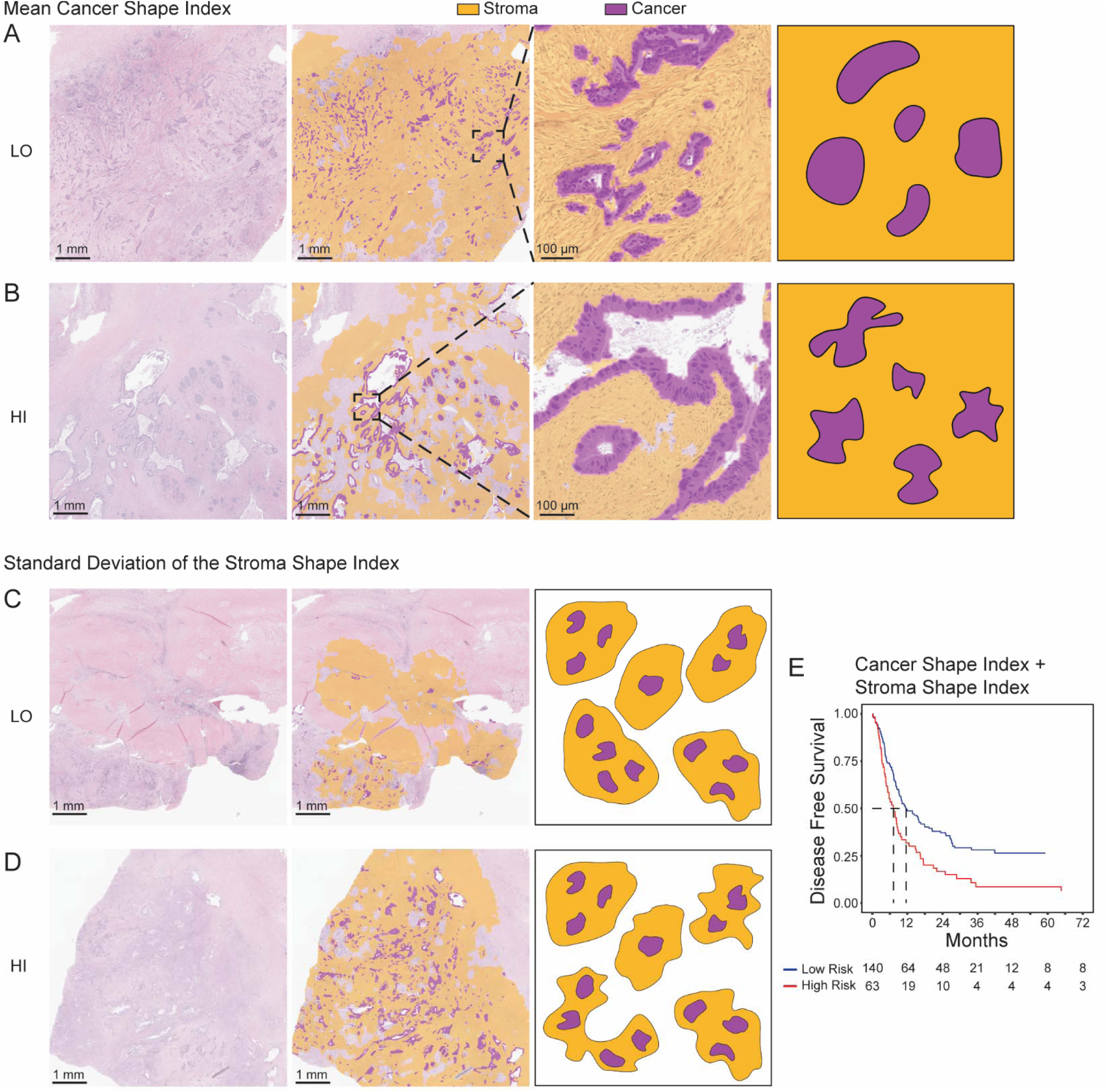
Shape index of residual cancer and stroma in post-treatment PDAC and associated DFS. (A, B) Representative cases illustrating high and low mean cancer shape index (CSI), respectively. (C, D) Representative cases illustrating high and low standard deviation of stroma shape index (SSI), respectively. Each panel (A–D) displays, from left to right: hematoxylin and eosin (H&E)–stained whole-slide images at 1× magnification, corresponding AI-generated overlays with cancer (purple) and stroma (yellow) segmentation, views at 10× magnification (for A, B), and a graphical representation of the spatial metric. Shape index (SI) is calculated as SI = P/(2√πA), where P is patch perimeter and A is patch area. A value of 1 indicates a perfectly circular shape, while higher values reflect increasing boundary irregularity. (E) Kaplan–Meier plot showing DFS stratified by risk group, defined by ln(mean CSI) and ln(SD SSI).

### Mean Stromal Area and Edge Density

An alternative statistical model employing backwards elimination (**Supplementary Table 5**) identified edge density (Figure 6A and B) and mean stroma area (Figure 6C and D) as significant spatial features. While the mean stroma area is self-explanatory, edge density quantifies the total length of all edge segments in a landscape, standardized by the total area of the landscape or patch, providing understanding of landscape fragmentation and complexity. Edge density is calculated as follows:

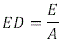

**Figure 6.**
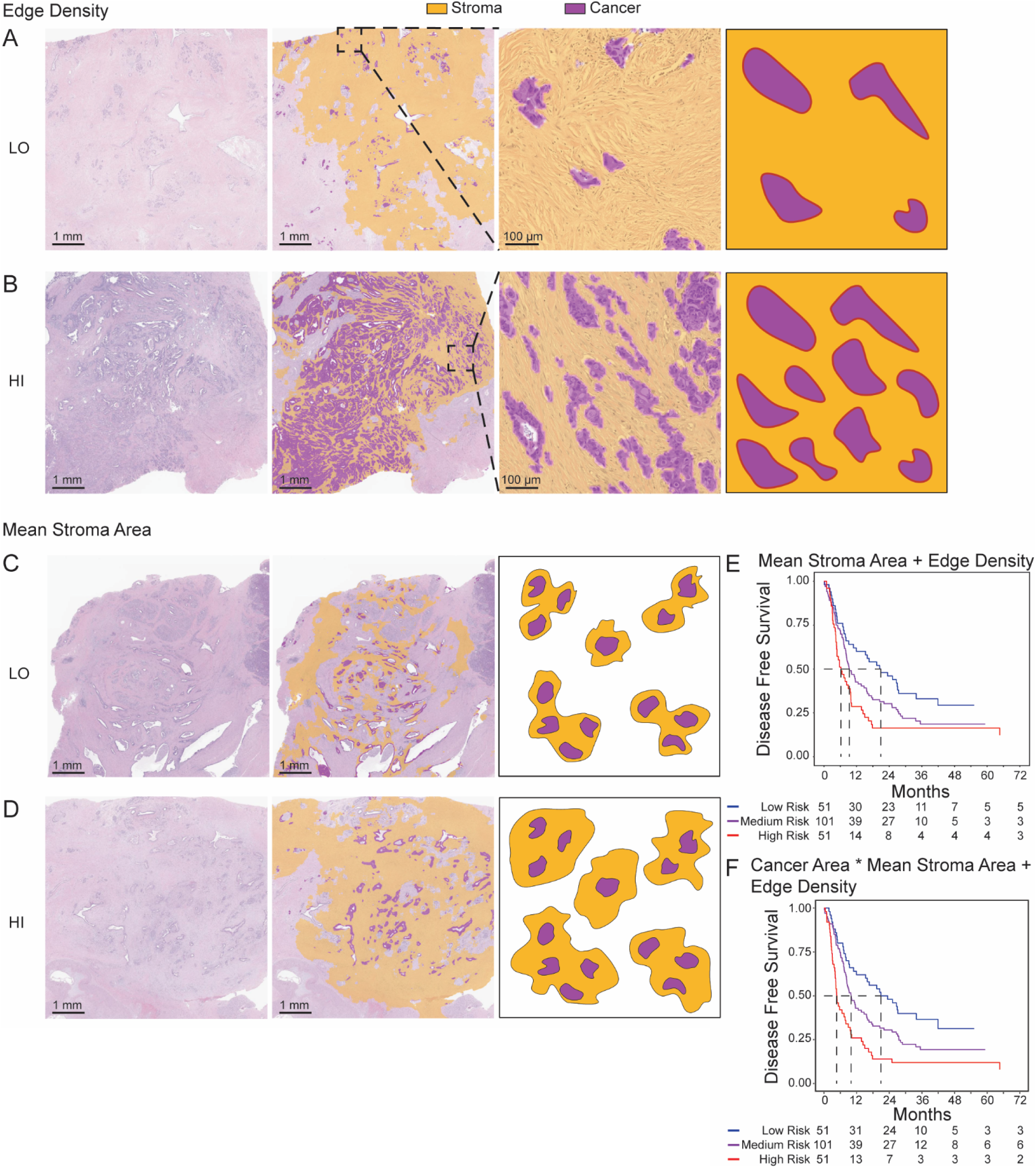
Edge density and mean stroma area in post-treatment PDAC and associated DFS. (A, B) Representative cases illustrating high and low edge density, respectively. (C, D) Representative cases illustrating high and low mean stroma area, respectively. Each panel (A–D) displays, from left to right: H&E–stained whole-slide images at 1× magnification, corresponding AI-generated overlays with cancer (purple) and stroma (yellow) segmentation, views at 10× magnification (for A, B), and a graphical representation of the spatial metric. (E) Kaplan–Meier plot showing DFS stratified by quartiles on a prognostic index derived from mean stroma area and edge density. (F) Prognostic refinement using an integrated model incorporating ln-transformed cancer area, men stroma area, and their interaction, demonstrating improved risk discrimination.

Where E represents the total length of edges and A is the total area of the landscape. Higher edge density values indicate greater fragmentation, while lower values are suggestive of a more compact, continuous habitat with fewer boundaries.

The prognostic index was calculated as 0.2701 * ln(mean stromal area) + 0.3905 * ln(edge density). High-risk patients (n=51) had a median DFS of 6.21 months (95% CI: 4.64 – 9.86), intermediate-risk patients (n=101) had a median DFS of 9.37 months (95% CI: 8.05 – 15.62), while low-risk patients (n=51) had a median DFS of 20.91 months (95% CI: 10.85 – 33.93) (Figure 6E). Multivariate Cox regression analysis confirmed that these spatial features are independent risk factors for DFS, with high-risk classification associated with a 1.94-fold higher risk of recurrence (HR=1.94, 95% CI: 1.21-3.10, p=0.006) (**Supplementary Table 6**). Sensitivity analysis also included other clinicopathologic covariates, which resulted in consistent results (**Supplementary Table 7**).

### Integration of Cancer Area

Extensive literature underscores that the extent of residual cancer, often subjectively assessed by tumor area or burden, holds independent clinical significance. To align our risk model with these historical insights, we incorporated the cancer area as an additional feature in the modeling process. In univariate analysis, the natural log-transformed cancer area demonstrated a p-value of 0.178 for association with DFS, meeting our threshold for variable advancement.

To evaluate the impact of integrating cancer area, we replaced edge density with ln(cancer area) in RPTR models, as well as in Cox proportional hazards models. Although RPTR models remained unchanged from prior analyses, backward selection approaches, which forced ln(cancer area) into the model, identified a prognostic combination of ln(cancer area), ln(mean stroma area), and ln(edge density).

Further refinement through interaction modeling revealed that the combination of ln(cancer area) and ln(mean stroma area), including their interaction term, significantly stratified patients into distinct risk categories for DFS (Figure 6F). This integrated model, adjusting for covariates, demonstrated improved discriminatory ability, with C-statistics reaching 0.666, supporting the additive prognostic value of cancer area alongside established spatial features. Collectively, these analyses validate the inclusion of cancer area in computational pathology-derived risk models, providing a framework that incorporates both tissue architecture and tumor burden as clinically important features in residual PDAC.

## Discussion

This study demonstrates that in PDAC treated with TNT and resection, but limited by minor pathologic response, the spatial architecture of the residual tumor-stroma ecosystem carries independent, clinically meaningful information about recurrence risk that is not captured by conventional response grading. Specifically, two multivariable spatial models including (1) a configuration model defined by mean CSI and variability of SSI and (2) a composition model defined by mean stromal area and landscape edge density, were independently associated with DFS after adjustment for established clinicopathologic covariates. Notably, neither CAP2 vs CAP3 response scoring, nor the area of residual cancer alone stratified DFS in this minor-response cohort, underscoring a cap in current assessment paradigms that our spatial approach begins to fill.

These results both refine and extend prior observations that major pathologic response is favorable in PDAC, whereas patients with residual disease show heterogeneous outcomes despite similar CAP grades^10–15^. Traditional grading emphasizes “how much” tumor remains and is vulnerable to inter-observer variability^12–15^. Our analysis asks a different question: how is the residual disease arranged? By quantifying patch shapes, sizes, densities, and the amount of cancer-stroma interface, we show that organization, not just extent, encodes treatment-resistant biology after TNT.

Viewed through an ecological lens^32^, several clear patterns emerged. First, greater fragmentation and mixing (higher edge density, higher diversity, and lower homotypic aggregation) was a hallmark of non-response at the cohort level and signaled worse DFS in multivariable models. Second, shape features mattered: lower mean CSI (more compact cancer patches) together with higher variability in SSI (more heterogeneous stromal shapes) defined as a high-risk configuration class. These observations align with mechanistic ideas that cancer–stroma interfaces act as privileged habitats for nutrient exchange^18–20^, immune evasion or exclusion ^21–25^, and mechanically favorable routes for invasion and collective migration^26–30^. An interface-rich, intermixed landscape creates more edge niches and a wider array of microenvironments for selection to exploit after therapy, enabling recurrence despite comparable disease burden.

The clinical message is pragmatic. First, post-TNT risk stratification improves when quantitative spatial metrics are added to standard pathology. Our CSI/SSI and stroma-area/edge-density models outperformed CAP grading in this setting and remained independent of ypT, ypN, margin status, and perineural or lymphovascular invasion. These signatures could help identify patients for intensified or mechanism-matched adjuvant strategies even when traditional indicators are equivocal. Second, the metrics are observer-independent and automatable on routine H&E slides, making them suitable for real-world deployment and for risk adaptive trials that already pivot on residual disease but currently lack spatial resolution^36–38^. For example, patients with high edge density and high stromal-shape variability might be prioritized for approaches that target edge biology (e.g., stroma-modulating agents, matrix/biomechanics pathways, or immunotherapies designed to access interface regions), whereas patients with coarser, more aggregated landscapes might warrant different strategies.

This work has limits. It is a retrospective, single-enterprise study in PDAC with minor regression. External validation is needed across scanners, staining protocols, and segmentation platforms. Although we applied quality controls, any AI-assisted pipeline can inherit biases from training data and class definitions. To maximize generalizability we focused on cancer vs stroma rather than finer cell states. Future studies should resolve cellular constituents at and across edges (e.g., fibroblast subtypes, immune phenotypes) to link architectural signatures to the actors and pathways that sustain persistence after therapy. Treatment-specific interactions and prospective integration into adjuvant decision workflows are important next steps.

Taken together, reframing treated PDAC as a post-therapy landscape reveals that the topology of residual cancer-stroma, not merely its quantity, tracks with therapeutic resistance and risk of relapse. Interface-rich fragmentation, compact cancer geometry, and heterogeneous stromal shapes define high-risk ecologies independent of standard features, whereas coarser, less interspersed architectures associate with longer DFS. These insights motivate spatially informed, risk adaptive adjuvant strategies and mechanistic studies of edge habitats and mixing as therapeutic vulnerabilities, with the broader aim of converting descriptive histology into actionable ecology for patients who need improved post-TNT outcomes

## Methods

### Study Design, Study Population, and Data Collection

All experiments in this study involving human tissue or data were conducted in accordance with the Declaration of Helsinki. Clinical data and tissue in the study were collected after Institutional Review Board review and approval (IRB 22-007646). All subjects had undergone surgical resection for PDAC within the Mayo Clinic Enterprise following TNT between 2010-2022. Variant exocrine carcinomas were excluded, in addition to those without research consent.

### H&E Slide Digitization

A digital pathology library was created, consisting of H&E stained WSI of surgically resected PDAC. The most representative slides from each case (an average of five slides per case, range of 1-16) were selected. Each slide was scanned at 40X magnification (0.25 microns/pixel) via high-throughput digital imaging by Aperio GT 450.

### PathExplore PDAC-Based Tissue Segmentation

To enable quantitative spatial analysis of the TME, we utilized PathExplore™ PDAC (v1.0), a commercially available artificial intelligence (AI)-driven platform developed by PathAI, Inc. PathExplore PDAC is designed for single-cell resolution spatial characterization of H&E-stained WSIs of PDAC specimens. It leverages deep learning models trained on pathologist-annotated datasets to identify and classify tissue compartments and cell types within the TME. PathExplore PDAC processed the digitized slides to perform automated tissue analysis, generating image overlays of cancer and cancer-associated stroma. Artifactual regions (e.g., folds, blur) were excluded via the PathExplore artifact model to ensure segmentation fidelity.

In this study, we specifically leveraged the spatial maps of cancer and stroma compartments to derive two-dimensional landscape mosaics. These served as the basis for TLA, a framework integrating landscape ecology metrics to quantify composition and configuration across the TME. PathExplore’s harmonized tissue classes enabled consistent segmentation across samples, ensuring robustness for downstream statistical modeling. The segmentation results formed the foundation for extracting patch– and landscape-level metrics, which were subsequently aggregated at the patient level for association with disease-free survival (DFS).

Model performance for cancer and stroma segmentation within PathExplore PDAC was benchmarked using established precision and recall thresholds to assess agreement with pathologist annotations. For regions where the model-predicted area of cancer or cancer-associated stroma exceeded 5% of usable tissue, segmentation quality was classified as “Excellent” if both precision and recall exceeded 91%, “Good” if both exceeded 80%, and “Notable Over– or Underestimation” based on imbalanced precision-recall profiles or discrepancies with ground truth annotations. For predicted areas below 5%, a modified framework prioritized high recall and absence of false positives to mitigate misclassification of rare features. Only segmentation outputs meeting “Excellent” or “Good” thresholds were included for landscape metric extraction and subsequent analyses, ensuring high-confidence tissue compartment delineation across the cohort (**Supplementary Figure 6**).

### Tumor Landscape Analysis Pipeline

Spatial metrics were computed using Tumor Landscape Analysis (TLA), a pipeline of analysis adapted from landscape ecology methodologies^39,40^ and recently described in detail.^32^ For spatial modeling using TLA, input for the pipeline was derived from cancer and cancer-associated stroma overlays, two-dimensional landscape mosaics, provided by PathExplore PDAC. Each mosaic consists of a two-dimensional raster array of discrete categorical labels describing distinct regional patches, with each label corresponding to a tissue compartment class. Analysis was restricted to the cancer and stromal components, as these features have been linked to long-term survival outcomes.^10,11^ Spatial metrics were computed at three hierarchical levels: the patch level, in which individual patches were quantified by their geometrical features, including area, edge length, and shape index; the landscape level, where whole-slide descriptors were generated based on the distributions of patch metrics and different measures of configuration and complexity of patch arrangements, including measures of diversity, entropy, and adjacency (interspersion); and the patient level, where measurements across multiple slides per patient were summarized by either aggregation or the mean, minimum, maximum and range.^20^

All code for data processing and spatial analysis associated with this study is publicly available in the CarrLab GitHub repository (https://github.com/CarrLab/TLA). Documentation for usage, updates, and versioning is maintained at this location.

### Outcomes Assessment

DFS was used as the primary endpoint of the study and is defined as the time from the date of curative resection to the date of first documented disease recurrence or death due to all causes, whichever occurred first.

### Statistical Analysis

Spatial pathological features (SPFs) were summarized by mean (standard of deviation [SD]), median, and range, compared by subgroups defined by clinical and disease characteristics. Two-level (high vs. low) risk classifications were developed based on the prognostic value of SPFs regarding DFs. Spearman correlation coefficients were used to describe the inter-correlations between SPFs to detect pairs with high collinearity. Normality transformations were performed when it is necessary to improve stability of the model fitting, by minimizing the influence of extreme values. Univariate association between individual SPF and DFS was assessed by Cox regression model, as well as spline fitting for non-linear effects. The individual SPFs with p-value < 0.20 were selected for subsequent modeling procedures for risk classification building. Two risk classification development methods were utilized. Recursive partitioning and tree regression method was used to split population into subgroups with homogeneity DFS which handles non-linear effects and interactions among SPFs naturally. High– vs. low-risk subgroups were defined by the end nodes generated by the tree method. The second method was based on traditional Cox regression with backward elimination procedure on selected SPFs after univariate analysis. The SPF variables with p value < 0.05 stayed in the final multivariable model. Interaction terms among remaining SPF variables were further evaluated. The final model was used to produce the continuous risk score for each patient. Since this score was associated with DFS in a linear form, median value was applied to define high-vs. low-risk groups. The discrimination abilities of both risk-classifications were quantified by C statistics. The risk-classifications were further evaluated by multivariable Cox models, adjusting for clinical and disease characteristics.

## Acknowledgements

We extend our deepest gratitude to our patients and their families. This work was also supported by the Gerstner Family Foundation Career Development Award, the Grand Forks Career Development Award, and the Mayo Clinic Center for Clinical and Translational Science (CCaTS) through a Small Grant (UL1TR000135). Ryan Carr gratefully acknowledges this funding support, which was instrumental in the development and execution of this study.

