## Supplemental Data for "Spatial Conformation of Pancreatic Cancer Is Associated with Disease Recurrence after Curative-Intent Total Neoadjuvant Therapy"

| **Suplementary Table 1. Univariate Associations Between Individual Clinical-pathological Factors and DFS** | | | | |
| --- | --- | --- | --- | --- |
| **Characteristic** | **N** | **HR***^1^* | **95% CI***^1^* | **p-value** |
| **Age at Surgery (5 year increments)** | 203 | 0.95 | 0.87, 1.04 | 0.25 |
| **Race** |  |  |  |  |
| White | 192 | — | — |  |
| Non-White | 9 | 0.81 | 0.36, 1.84 | 0.62 |
| **Gender** |  |  |  |  |
| Male | 107 | — | — |  |
| Female | 96 | 1.04 | 0.76, 1.43 | 0.79 |
| **Clinical Stage at Diagnosis** |  |  |  |  |
| IA | 23 | — | — |  |
| Cannot Be Determined | 1 | 1.15 | 0.15, 8.64 | 0.89 |
| IB | 61 | 0.75 | 0.43, 1.30 | 0.30 |
| IIA | 12 | 1.18 | 0.54, 2.57 | 0.67 |
| IIB | 51 | 0.88 | 0.50, 1.55 | 0.66 |
| III | 52 | 1.35 | 0.78, 2.35 | 0.28 |
| **Type of Surgical Resection** |  |  |  |  |
| Total Pancreatectomy | 32 | — | — |  |
| Distal Pancreatectomy | 45 | 0.94 | 0.56, 1.55 | 0.80 |
| Pancreaticoduodenectomy | 126 | 0.70 | 0.45, 1.08 | 0.11 |
| **Time from Diagnosis to Surgery (3 month increments)** | 203 | 1.01 | 0.87, 1.19 | 0.86 |
| **Histological Grade** |  |  |  |  |
| Moderately or Well-Differentiated | 142 | — | — |  |
| Undifferentiated, Poorly Differentiated, Or Cannot be Determined | 61 | 0.91 | 0.64, 1.29 | 0.59 |
| **Local Invasion** |  |  |  |  |
| Confined to the Pancreas | 54 | --- | --- |  |
| Other | 149 | 1.42 | 0.98, 2.05 | 0.062 |
| **Pathologic Response (CAP)** |  |  |  |  |
| No Response | 34 | — | — |  |
| Complete or Near Complete Response | 2 | 0.00 | 0.00, Inf | >0.99 |
| Partial Response | 154 | 0.67 | 0.44, 1.00 | 0.051 |
| **Resection Outcome** |  |  |  |  |
| R0 | 174 | — | — |  |
| R1 | 27 | 1.28 | 0.82, 2.02 | 0.28 |
| R2 | 1 | 16.7 | 2.17, 128 | **0.007** |
| **Lymphovascular Invasion** |  |  |  |  |
| No | 159 | — | — |  |
| Yes | 27 | 1.59 | 1.00, 2.51 | **0.048** |
| Indeterminate | 2 | 0.28 | 0.04, 2.07 | 0.21 |
| **Perineural Invasion** |  |  |  |  |
| No | 80 | — | — |  |
| Yes | 110 | 1.38 | 0.98, 1.93 | 0.063 |
| Indeterminate | 2 | 0.32 | 0.04, 2.44 | 0.27 |
| **Involved Margins at Resection** |  |  |  |  |
| No | 176 | — | — |  |
| Yes | 27 | 1.27 | 0.81, 2.00 | 0.29 |
| **Pathologic T Stage** |  |  |  |  |
| 1 | 46 | — | — |  |
| 2 | 102 | 1.36 | 0.90, 2.07 | 0.15 |
| 3 | 44 | 2.18 | 1.36, 3.48 | **0.001** |
| 4 | 10 | 3.46 | 1.63, 7.34 | **0.001** |
| **Pathologic N Stage** |  |  |  |  |
| Negative | 140 | --- | --- |  |
| Positive | 63 | 1.71 | 1.22, 2.38 | **0.002** |
| *^1^*HR = Hazard Ratio, CI = Confidence Interval | | | | |

**Supplementary Figure 1.** Correlation between spatial features and histologic grade. ns=not significant

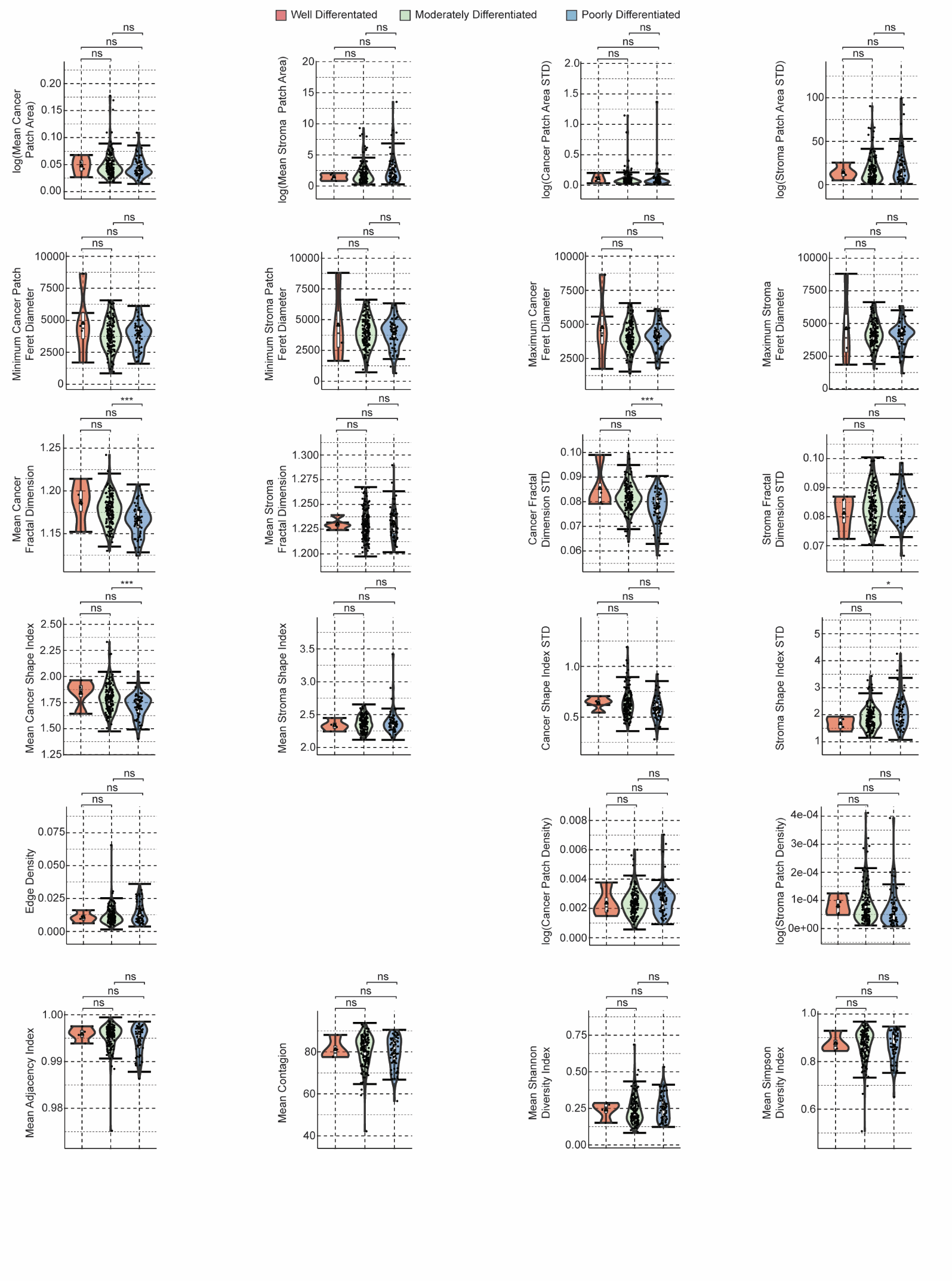

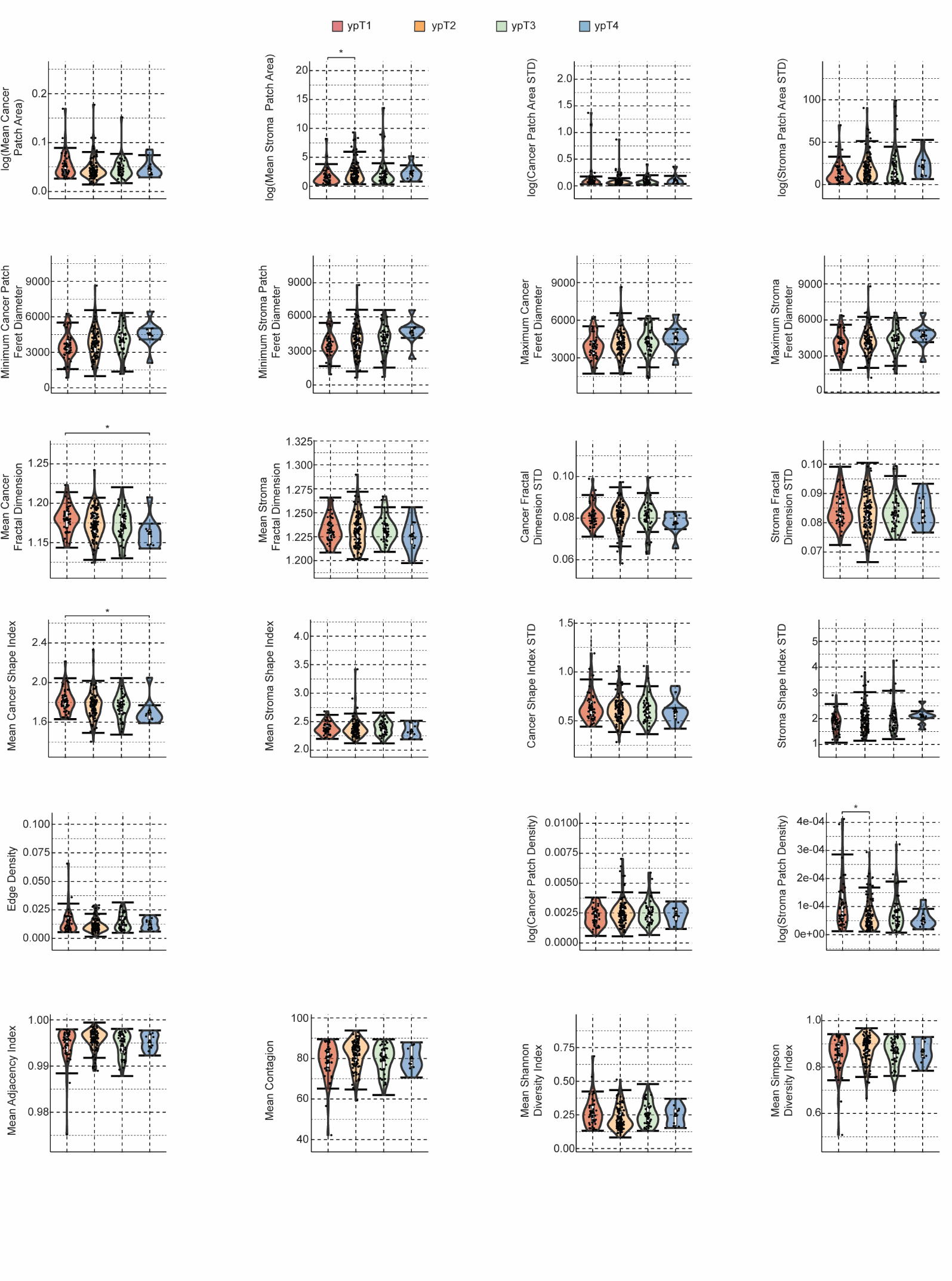

**Supplementary Figure 2.** Correlation between spatial features and pathologic tumor score (ypT). Only statistically significant differences are annotated.

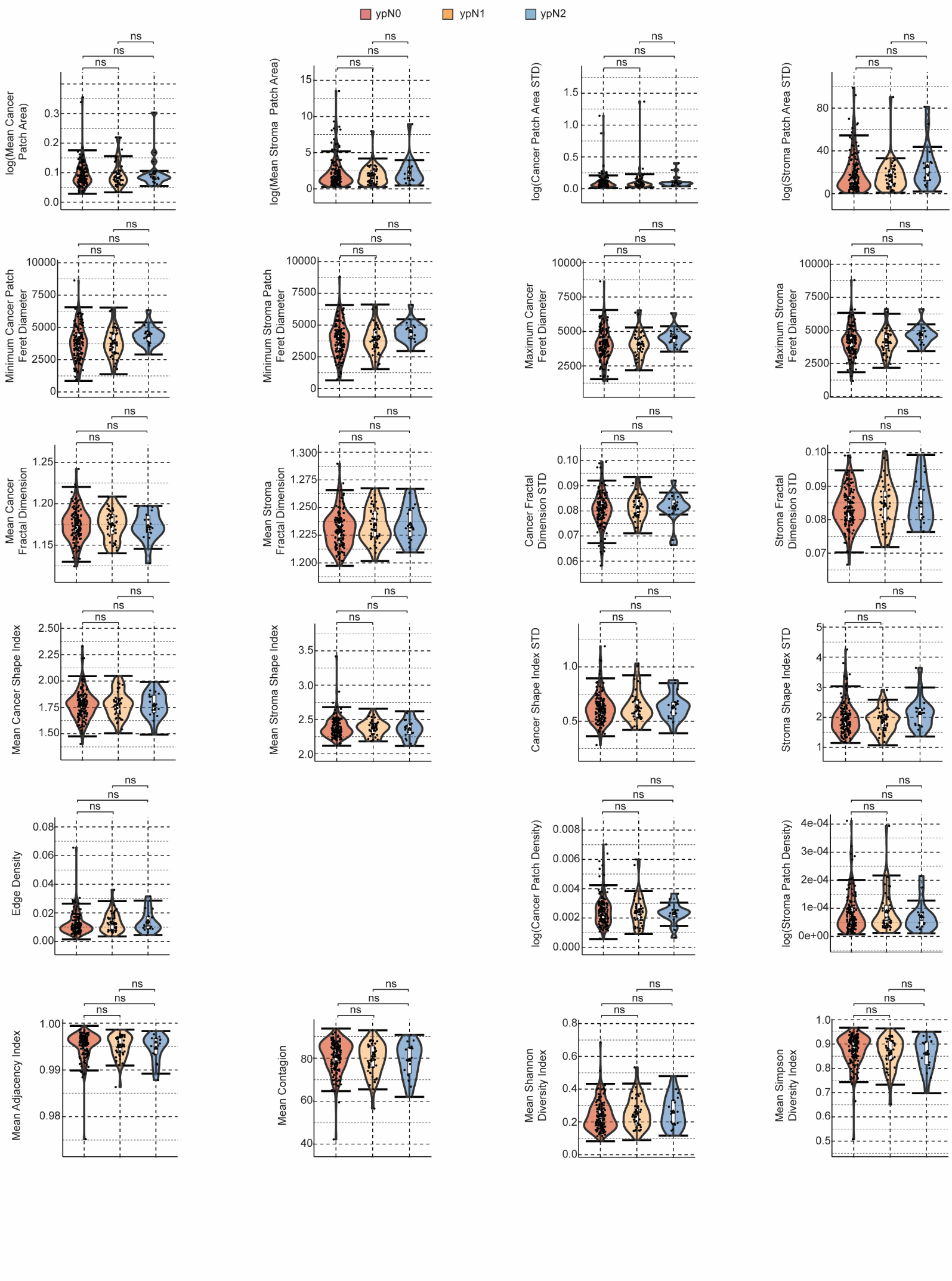

**Supplementary Figure 3.** Correlation between spatial features and pathologic nodal status (ypN). ns = not significant

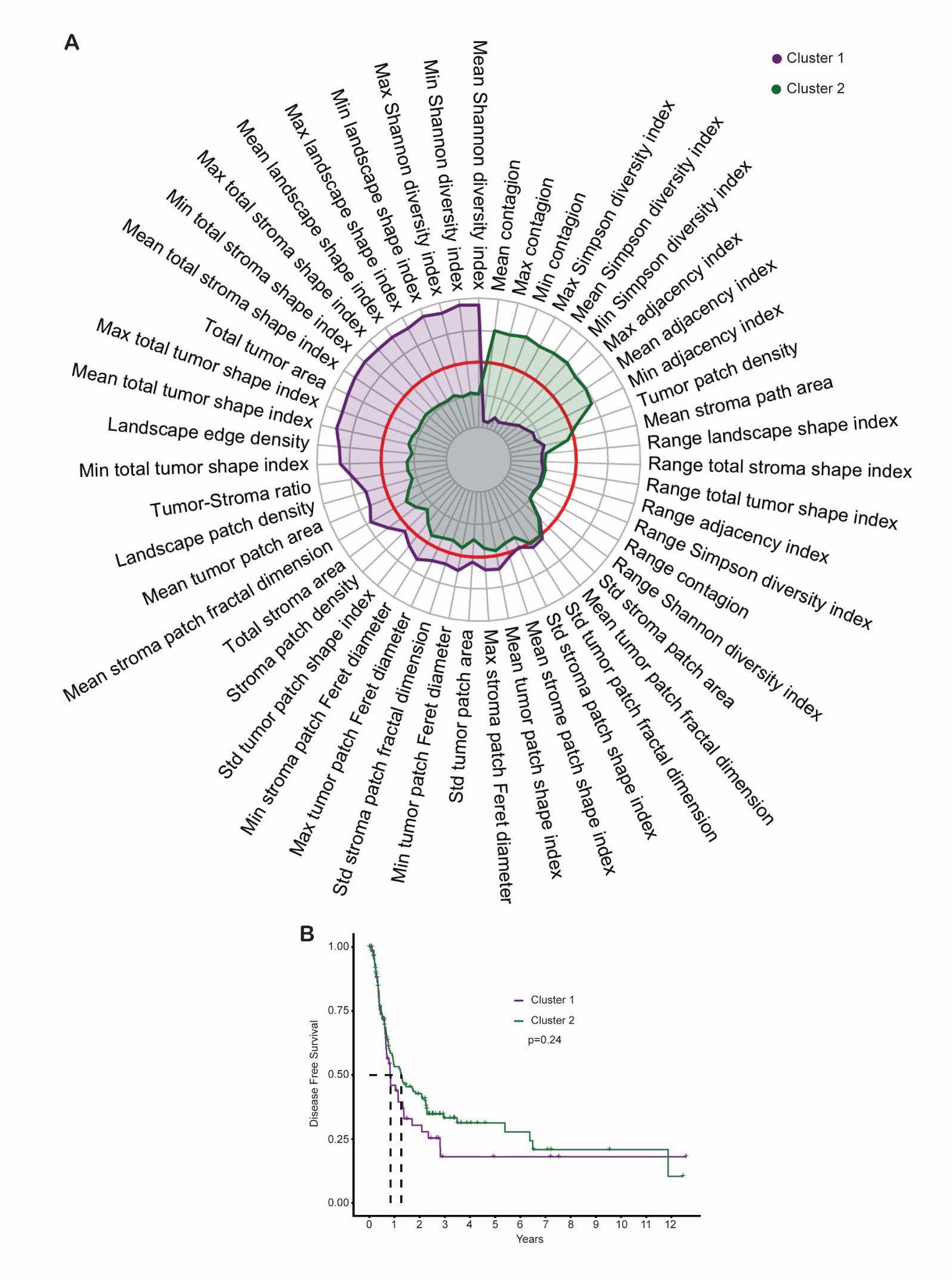

**Supplementary Figure 4.** A. Radar plot for median values of normalized spatial pathology features in each cluster of patients (as presented in Figure 4). The red circle indicates a zero value. Therefore, the median values have a clear separation between cluster for features in the upper two quadrats of the plot. B. Kaplan Meier curve for the two clusters of patients, indicating that, though some spatial pathology features are clearly distinct between patient groups, they are not predictive of outcomes, most likely due to all the features that overlap between groups (lower side of radar plot).

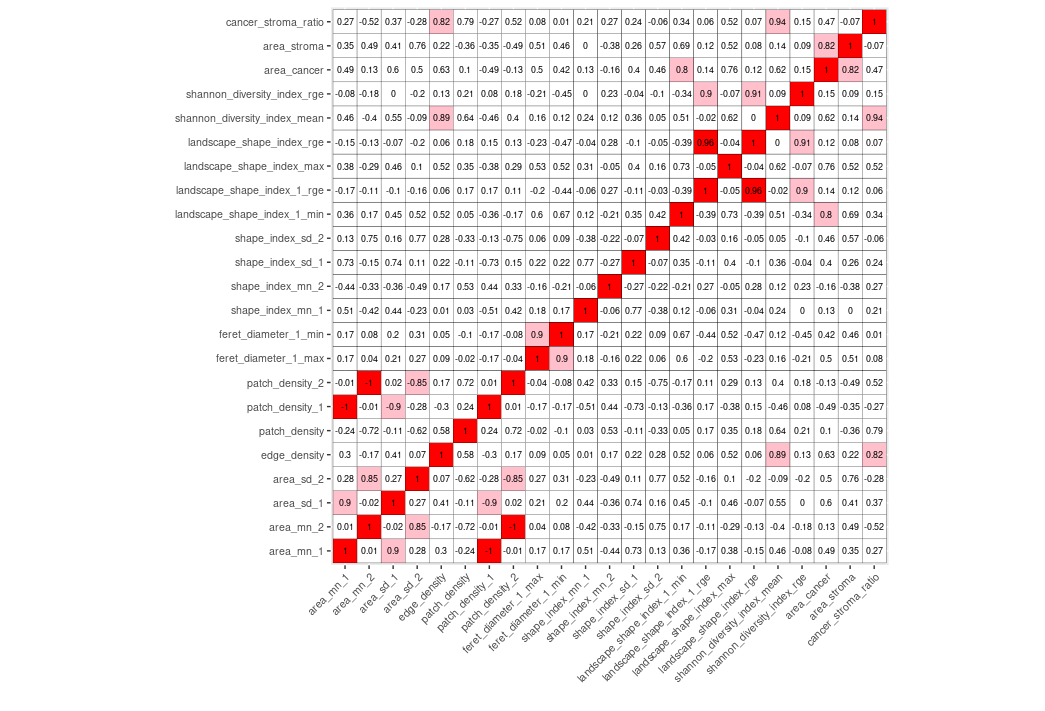

**Supplementary Figure 5.** Correlation matrix of all spatial features (Spearman Rho)

| **Supplementary Table 2. Demographics and Clinicopathological Factors per Cancer and Stroma Shape Index Risk Model** | | | | | | | | |
| --- | --- | --- | --- | --- | --- | --- | --- | --- |
|  | **High Risk (N=63)** | | | **Low Risk (N=140)** | | **Total (N=203)** | | **p value** |
| **Age at Surgery** | |  |  | |  | | **0.6164 ^(1)^** | |
| Mean (SD) | | 65.9 (8.3) | 65.1 (9.6) | | 65.4 (9.2) | |  | |
| Median (Range) | | 66.8 (47.0, 80.5) | 66.2 (39.4, 85.3) | | 66.5 (39.4, 85.3) | |  | |
| **Race** | |  |  | |  | | **0.7240 ^(2)^** | |
| White | | 60 (96.8%) | 132 (95.0%) | | 192 (95.5%) | |  | |
| Non-White | | 2 (3.2%) | 7 (5.0%) | | 9 (4.5%) | |  | |
| **Sex** | |  |  | |  | | **0.1453 ^(3)^** | |
| Female | | 25 (39.7%) | 71 (50.7%) | | 96 (47.3%) | |  | |
| Male | | 38 (60.3%) | 69 (49.3%) | | 107 (52.7%) | |  | |
| **Clinical Stage at Diagnosis** | |  |  | |  | | **0.5969 ^(2)^** | |
| Cannot Be Determined | | 0 (0.0%) | 1 (0.7%) | | 1 (0.5%) | |  | |
| IA | | 6 (9.8%) | 17 (12.2%) | | 23 (11.5%) | |  | |
| IB | | 16 (26.2%) | 45 (32.4%) | | 61 (30.5%) | |  | |
| IIA | | 4 (6.6%) | 8 (5.8%) | | 12 (6.0%) | |  | |
| IIB | | 14 (23.0%) | 37 (26.6%) | | 51 (25.5%) | |  | |
| III | | 21 (34.4%) | 31 (22.3%) | | 52 (26.0%) | |  | |
| **Type of Surgical Resection** | |  |  | |  | | **0.1575 ^(3)^** | |
| Distal Pancreatectomy | | 18 (28.6%) | 27 (19.3%) | | 45 (22.2%) | |  | |
| Pancreaticoduodenectomy | | 33 (52.4%) | 93 (66.4%) | | 126 (62.1%) | |  | |
| Total Pancreatectomy | | 12 (19.0%) | 20 (14.3%) | | 32 (15.8%) | |  | |
| **Time from Diagnosis to Surgery (months)** | |  |  | |  | | **0.4500 ^(1)^** | |
| Mean (SD) | | 8.3 (3.6) | 8.7 (3.0) | | 8.6 (3.2) | |  | |
| Median (Range) | | 8.0 (1.8, 18.2) | 8.2 (2.9, 19.4) | | 8.2 (1.8, 19.4) | |  | |
| **Histological Grade** | |  |  | |  | | **<0.0001 ^(2)^** | |
| 1 - Well-differentiated | | 0 (0.0%) | 4 (2.9%) | | 4 (2.0%) | |  | |
| 2 - Moderately differentiated | | 32 (50.8%) | 106 (75.7%) | | 138 (68.0%) | |  | |
| 3 - Poorly differentiated | | 29 (46.0%) | 28 (20.0%) | | 57 (28.1%) | |  | |
| 4 - Undifferentiated | | 2 (3.2%) | 0 (0.0%) | | 2 (1.0%) | |  | |
| X - Cannot be determined | | 0 (0.0%) | 2 (1.4%) | | 2 (1.0%) | |  | |
| **Local Invasion** | |  |  | |  | | **0.0317 ^(2)^** | |
| Confined to Pancreas | | 10 (16.9%) | 44 (31.9%) | | 54 (27.4%) | |  | |
| Duodenal Wall | | 3 (5.1%) | 20 (14.5%) | | 23 (11.7%) | |  | |
| Peripancreatic Soft Tissues | | 24 (40.7%) | 40 (29.0%) | | 64 (32.5%) | |  | |
| Multiple | | 18 (30.5%) | 29 (21.0%) | | 47 (23.9%) | |  | |
| Other | | 3 (5.1%) | 4 (2.9%) | | 7 (3.6%) | |  | |
| None | | 1 (1.7%) | 1 (0.7%) | | 2 (1.0%) | |  | |
| **Pathologic Response** (CAP) | |  |  | |  | | **0.2782 ^(2)^** | |
| 2 - Partial Response | | 42 (76.4%) | 114 (84.5%) | | 156 (82.2%) | |  | |
| 3 - No Response | | 13 (23.6%) | 21 (15.6%) | | 34 (17.9%) | |  | |
| **Resection Outcome** | |  |  | |  | | **0.3806 ^(2)^** | |
| R0 | | 53 (84.1%) | 121 (87.1%) | | 174 (86.1%) | |  | |
| R1 | | 9 (14.3%) | 18 (12.9%) | | 27 (13.4%) | |  | |
| R2 | | 1 (1.6%) | 0 (0.0%) | | 1 (0.5%) | |  | |
| **Lymphovascular Invasion** | |  |  | |  | | **0.2786 ^(2)^** | |
| Yes | | 11 (20.0%) | 16 (12.0%) | | 27 (14.4%) | |  | |
| No | | 44 (80.0%) | 115 (86.5%) | | 159 (84.6%) | |  | |
| **Perineural Invasion** | |  |  | |  | | **0.0009 ^(2)^** | |
| Yes | | 43 (76.8%) | 67 (49.3%) | | 110 (57.3%) | |  | |
| No | | 13 (23.2%) | 67 (49.3%) | | 80 (41.7%) | |  | |
| **Pathologic T Stage** | |  |  | |  | | **0.0425 ^(2)^** | |
| 1a | | 0 (0.0%) | 7 (5.0%) | | 7 (3.5%) | |  | |
| 1b | | 1 (1.6%) | 3 (2.2%) | | 4 (2.0%) | |  | |
| 1c | | 8 (12.7%) | 27 (19.4%) | | 35 (17.3%) | |  | |
| 2 | | 34 (54.0%) | 68 (48.9%) | | 102 (50.5%) | |  | |
| 3 | | 13 (20.6%) | 31 (22.3%) | | 44 (21.8%) | |  | |
| 4 | | 7 (11.1%) | 3 (2.2%) | | 10 (5.0%) | |  | |
| **Number of Nodes Removed** | |  |  | |  | | **0.3147 ^(1)^** | |
| Median (Range) | | 19.0 (1.0, 56.0) | 20.0 (3.0, 71.0) | | 20.0 (1.0, 71.0) | |  | |
| **Number of Positive Nodes** | |  |  | |  | | **0.8709 ^(1)^** | |
| Median (Range) | | 0.0 (0.0, 10.0) | 0.0 (0.0, 17.0) | | 0.0 (0.0, 17.0) | |  | |
| **Pathological N Stage** | |  |  | |  | | **0.6281 ^(3)^** | |
| N0 (0 LN+) | | 44 (69.8%) | 96 (68.6%) | | 140 (69.0%) | |  | |
| N1 (1-3 LN+) | | 12 (19.0%) | 33 (23.6%) | | 45 (22.2%) | |  | |
| N2 (4+ LN+) | | 7 (11.1%) | 11 (7.9%) | | 18 (8.9%) | |  | |
| **Neoadjuvant Mode of Treatment** | |  |  | |  | | **0.0985 ^(3)^** | |
| Chemo and RT with radiosensitizing chemo | | 42 (66.7%) | 107 (76.4%) | | 149 (73.4%) | |  | |
| Chemo and RT w/o radiosensitizing chemo | | 1 (1.6%) | 1 (0.7%) | | 2 (1.0%) | |  | |
| Chemo only | | 14 (22.2%) | 29 (20.7%) | | 43 (21.2%) | |  | |
| RT with radiosensitizing chemo | | 6 (9.5%) | 3 (2.1%) | | 9 (4.4%) | |  | |

1. Wilcoxon rank sum test; 2. Fisher’s Exact Test for Count Data; 3. Pearson’s Chi-squared test

| **Supplementary Table 3. Multivariable Cox Analysis (DFS) Cancer and Stroma Shape Index Risk Model** | | | |
| --- | --- | --- | --- |
| **Characteristic** | **HR***^1^* | **95% CI***^1^* | **p-value** |
| **Risk Category** | | | |
| Low Risk | — | — |  |
| High Risk | 1.64 | 1.18, 2.30 | **0.004** |
| **Pathologic T Stage** |  |  |  |
| 1 | — | — |  |
| 2 | 1.25 | 0.82, 1.92 | 0.294 |
| 3 | 2.06 | 1.27, 3.33 | **0.003** |
| 4 | 2.89 | 1.35, 6.19 | **0.006** |
| **Pathologic Nodal Involvement** |  |  |  |
| Negative | — | — |  |
| Positive | 1.82 | 1.29, 2.55 | **<0.001** |
| **c-index** = 0.628; Log-likelihood = -698; No. Obs. = 202; N events = 157 | | | |
| *^1^*HR = Hazard Ratio, CI = Confidence Interval | | | |

| **Supplementary Table 4. Expanded Multivariable Cox Analysis (DFS) Cancer and Stroma Shape Index Risk Model** | | | |
| --- | --- | --- | --- |
| **Characteristic** | **HR***^1^* | **95% CI***^1^* | **p-value** |
| **Risk Category** |  |  |  |
| Low Risk | — | — |  |
| High Risk | 1.75 | 1.17, 2.63 | **0.007** |
| **Lymphovascular Invasion** |  |  |  |
| No | — | — |  |
| Yes | 1.22 | 0.71, 2.08 | 0.471 |
| Indeterminate | 0.43 | 0.06, 3.39 | 0.425 |
| **Resection Outcome** |  |  |  |
| R0 | — | — |  |
| R1 | 0.91 | 0.56, 1.48 | 0.701 |
| R2 | 6.50 | 0.74, 56.8 | **0.090** |
| **T-Stage (Pathologic)** |  |  |  |
| 1 | — | — |  |
| 2 | 1.09 | 0.67, 1.77 | 0.724 |
| 3 | 2.12 | 1.20, 3.73 | **0.010** |
| 4 | 2.26 | 0.93, 5.47 | **0.072** |
| **Node Positive (Pathologic)** |  |  |  |
| Negative | — | — |  |
| Positive | 1.73 | 1.19, 2.52 | **0.004** |
| **Histologic Grade** |  |  |  |
| Moderately or Well-Differentiated | — | — |  |
| Undifferentiated, Poorly Differentiated, Or Cannot be Determined | 0.70 | 0.46, 1.07 | 0.100 |
| **Local Invasion** |  |  |  |
| Confined to Pancreas | — | — |  |
| Other | 1.14 | 0.74, 1.76 | 0.541 |
| **Perineural Invasion** |  |  |  |
| No | — | — |  |
| Yes | 1.04 | 0.71, 1.54 | 0.837 |
| Indeterminate |  |  |  |
| *^1^*HR = Hazard Ratio, CI = Confidence Interval | | | |
| **c-index = 0.627;** Log-likelihood = -632; No. Obs. = 188; N events = 144 | | | |

**Supplementary Table 5. Demographics and Clinicopathological Factors per Stroma Area + Edge Density Risk Model**

|  | **Low Risk (25%) (N=51)** | **Middle 50% (N=101)** | **High Risk (25%) (N=51)** | **Total (N=203)** | **p value** |
| --- | --- | --- | --- | --- | --- |
| **Age at Surgery** |  |  |  |  | 0.063 |
| Mean (SD) | 65.2 (9.7) | 64.3 (9.1) | 67.8 (8.8) | 65.4 (9.2) |  |
| Median (Range) | 66.1 (45.7, 85.3) | 65.0 (39.4, 83.4) | 69.4 (47.0, 80.5) | 66.5 (39.4, 85.3) |  |
| **Race** |  |  |  |  | 0.744 |
| Non-White | 3 (5.9%) | 5 (5.0%) | 1 (2.0%) | 9 (4.5%) |  |
| White | 48 (94.1%) | 96 (95.0%) | 48 (98.0%) | 192 (95.5%) |  |
| **Gender** |  |  |  |  | 0.147 |
| Female | 26 (51.0%) | 52 (51.5%) | 18 (35.3%) | 96 (47.3%) |  |
| Male | 25 (49.0%) | 49 (48.5%) | 33 (64.7%) | 107 (52.7%) |  |
| **Clinical Stage at Diagnosis** |  |  |  |  | 0.334 |
| Cannot Be Determined | 0 (0.0%) | 1 (1.0%) | 0 (0.0%) | 1 (0.5%) |  |
| IA | 8 (15.7%) | 8 (8.0%) | 7 (14.3%) | 23 (11.5%) |  |
| IB | 14 (27.5%) | 29 (29.0%) | 18 (36.7%) | 61 (30.5%) |  |
| IIA | 0 (0.0%) | 8 (8.0%) | 4 (8.2%) | 12 (6.0%) |  |
| IIB | 17 (33.3%) | 26 (26.0%) | 8 (16.3%) | 51 (25.5%) |  |
| III | 12 (23.5%) | 28 (28.0%) | 12 (24.5%) | 52 (26.0%) |  |
| **Type of Surgical Resection** |  |  |  |  | 0.524 |
| Distal Pancreatectomy | 10 (19.6%) | 21 (20.8%) | 14 (27.5%) | 45 (22.2%) |  |
| Pancreaticoduodenectomy | 35 (68.6%) | 60 (59.4%) | 31 (60.8%) | 126 (62.1%) |  |
| Total Pancreatecomy | 6 (11.8%) | 20 (19.8%) | 6 (11.8%) | 32 (15.8%) |  |
| **Time from Diagnosis to Surgery (months)** |  |  |  |  | 0.087 |
| Mean (SD) | 8.8 (3.2) | 8.9 (3.1) | 7.6 (3.2) | 8.6 (3.2) |  |
| Median (Range) | 8.2 (3.5, 18.8) | 8.5 (3.5, 19.4) | 7.5 (1.8, 14.3) | 8.2 (1.8, 19.4) |  |
| **Histological Grade** |  |  |  |  | 0.05 |
| 1 - Well-differentiated | 1 (2.0%) | 2 (2.0%) | 1 (2.0%) | 4 (2.0%) |  |
| 2 - Moderately differentiated | 39 (76.5%) | 73 (72.3%) | 26 (51.0%) | 138 (68.0%) |  |
| 3 - Poorly differentiated | 10 (19.6%) | 24 (23.8%) | 23 (45.1%) | 57 (28.1%) |  |
| 4 – Undifferentiated | 0 (0.0%) | 1 (1.0%) | 1 (2.0%) | 2 (1.0%) |  |
| X - Cannot be determined | 1 (2.0%) | 1 (1.0%) | 0 (0.0%) | 2 (1.0%) |  |
| **Local Invasion** |  |  |  |  | 0.046 |
| Confined to Pancreas | 20 (39.2%) | 25 (24.8%) | 9 (17.6%) | 54 (26.6%) |  |
| Other | 31 (60.8%) | 76 (75.2%) | 42 (82.4%) | 149 (73.4%) |  |
| **Major Pathologic Response** |  |  |  |  | < 0.001 |
| Complete or Near Complete Response | 2 (4.1%) | 0 (0.0%) | 0 (0.0%) | 2 (1.1%) |  |
| No Response | 1 (2.0%) | 14 (14.4%) | 19 (43.2%) | 34 (17.9%) |  |
| Partial Response | 46 (93.9%) | 83 (85.6%) | 25 (56.8%) | 154 (81.1%) |  |
| **Resection Outcome** |  |  |  |  | 0.158 |
| R0 | 46 (92.0%) | 85 (84.2%) | 43 (84.3%) | 174 (86.1%) |  |
| R1 | 3 (6.0%) | 16 (15.8%) | 8 (15.7%) | 27 (13.4%) |  |
| R2 | 1 (2.0%) | 0 (0.0%) | 0 (0.0%) | 1 (0.5%) |  |
| **Lymphovascular Invasion** |  |  |  |  | 0.217 |
| Yes | 3 (6.4%) | 15 (15.3%) | 9 (20.9%) | 27 (14.4%) |  |
| No | 43 (91.5%) | 82 (83.7%) | 34 (79.1%) | 159 (84.6%) |  |
| Indeterminate | 1 (2.1%) | 1 (1.0%) | 0 (0.0%) | 2 (1.1%) |  |
| **Perineural Invasion** |  |  |  |  | < 0.001 |
| Yes | 19 (38.8%) | 55 (55.0%) | 36 (83.7%) | 110 (57.3%) |  |
| No | 29 (59.2%) | 44 (44.0%) | 7 (16.3%) | 80 (41.7%) |  |
| Indeterminate | 1 (2.0%) | 1 (1.0%) | 0 (0.0%) | 2 (1.0%) |  |
| **Involved Margins at Resection** |  |  |  |  | 0.201 |
| No | 48 (94.1%) | 85 (84.2%) | 43 (84.3%) | 176 (86.7%) |  |
| Yes | 3 (5.9%) | 16 (15.8%) | 8 (15.7%) | 27 (13.3%) |  |
| **Pathologic T Stage** |  |  |  |  | 0.223 |
| 1 | 16 (32.0%) | 19 (18.8%) | 11 (21.6%) | 46 (22.8%) |  |
| 2 | 24 (48.0%) | 54 (53.5%) | 24 (47.1%) | 102 (50.5%) |  |
| 3 | 10 (20.0%) | 20 (19.8%) | 14 (27.5%) | 44 (21.8%) |  |
| 4 | 0 (0.0%) | 8 (7.9%) | 2 (3.9%) | 10 (5.0%) |  |
| **Number of Nodes Removed** |  |  |  |  | 0.338 |
| Median (Range) | 21.0 (3.0, 40.0) | 19.0 (1.0, 71.0) | 17.0 (3.0, 56.0) | 20.0 (1.0, 71.0) |  |
| **Number of Positive Nodes** |  |  |  |  | 0.251 |
| Median (Range) | 0.0 (0.0, 4.0) | 0.0 (0.0, 13.0) | 0.0 (0.0, 17.0) | 0.0 (0.0, 17.0) |  |
| **Pathologic N Stage** |  |  |  |  | 0.399 |
| Negative | 39 (76.5%) | 66 (65.3%) | 35 (68.6%) | 140 (69.0%) |  |
| Positive | 12 (23.5%) | 35 (34.7%) | 16 (31.4%) | 63 (31.0%) |  |
| **Neoadjuvant Mode of Treatment** |  |  |  |  | 0.004 |
| Chemo and Radiation Regimens with Chemosensitizing | 41 (80.4%) | 81 (80.2%) | 27 (52.9%) | 149 (73.4%) |  |
| Chemo and Radiation Regimens without Chemosensitizing | 0 (0.0%) | 2 (2.0%) | 0 (0.0%) | 2 (1.0%) |  |
| Chemo Only | 8 (15.7%) | 15 (14.9%) | 20 (39.2%) | 43 (21.2%) |  |
| Radiation Regimen with Chemosensitizing | 2 (3.9%) | 3 (3.0%) | 4 (7.8%) | 9 (4.4%) |  |

| **Table 6. Multivariable Cox Analysis (DFS) Stroma Area + Edge Density Risk Model** | | | |
| --- | --- | --- | --- |
| **Characteristic** | **HR***^1^* | **95% CI***^1^* | **p-value** |
| **Risk Category** \| \| \|; Low Risk \| — \| — \|; Middle Risk \| 1.30 \| 0.86, 1.98 \| 0.200; High Risk \| 1.86 \| 1.17, 2.96 \| **0.009**; **Pathologic T Stage** \| \| \|; 1 \| — \| — \|; 2 \| 1.30 \| 0.85, 1.98 \| 0.200; 3 \| 2.01 \| 1.24, 3.25 \| **0.005**; 4 \| 3.16 \| 1.47, 6.82 \| **0.003**; **Pathologic Nodal Involvement** \| \| \|; Negative \| — \| — \|; Positive \| 1.77 \| 1.26, 2.48 \| **0.001**; **c-index** = 0.633; Log-likelihood = -699; No. Obs. = 202; N events = 157 \| \| \| | | | |
| *^1^*HR = Hazard Ratio, CI = Confidence Interval | | | |

| **Supplementary Table 7. Multivariable Cox Analysis (DFS) Stroma Area + Edge Density Risk Model** | | |  |
| --- | --- | --- | --- |
|  | **HR** | **95% CI** | **p-value** |
| **Risk Category** |  |  |  |
| Low Risk (25%) | — | — |  |
| Middle 50% | 1.27 | 0.81, 1.99 | 0.291 |
| High Risk (25%) | 1.94 | 1.10, 3.43 | 0.023 |
| **Lymphovascular Invasion** |  |  |  |
| No | — | — |  |
| Yes | 1.08 | 0.63, 1.85 | 0.788 |
| Indeterminate | 0.48 | 0.06, 3.57 | 0.474 |
| **Resection Outcome** |  |  |  |
| R0 | — | — |  |
| R1 | 0.99 | 0.60, 1.64 | 0.98 |
| R2 | 19 | 2.12, 170 | 0.009 |
| **T-Stage (Pathologic)** |  |  |  |
| 1 | — | — |  |
| 2 | 1.13 | 0.71, 1.80 | 0.595 |
| 3 | 1.93 | 1.13, 3.28 | 0.015 |
| 4 | 2.18 | 0.89, 5.33 | 0.086 |
| **Node Positive (Pathologic)** |  |  |  |
| Negative | — | — |  |
| Positive | 1.64 | 1.12, 2.40 | 0.011 |
| **Histologic Grade** |  |  |  |
| Moderately or Well-Differentiated | — | — |  |
| Undifferentiated, Poorly Differentiated, Or Cannot be Determined | 0.89 | 0.58, 1.36 | 0.579 |
| **Major Pathologic Response** |  |  |  |
| No Response | — | — |  |
| Near Complete Response | 0 | 0.00, Inf | 0.994 |
| Partial Response | 0.8 | 0.50, 1.26 | 0.333 |
| **Perineural Invasion** |  |  |  |
| No | — | — |  |
| Yes | 1.09 | 0.74, 1.59 | 0.675 |
| Indeterminate |  |  |  |
| **Number of Nodes Removed** | 1.02 | 1.00, 1.03 | 0.07 |
| Abbreviations: CI = Confidence Interval, HR = Hazard Ratio | | | |
| c-index = 0.656; Log-likelihood = -612; No. Obs. = 184; N events = 140 | | | |

**Supplementary Figure 6. Performance of the PathExplore PDAC model for tissue segmentation.** The model’s ability to identify cancer and cancer-associated stroma in PDAC was evaluated by comparing its predictions to expert pathologist annotations. Segmentation was rated “Excellent” if both precision and recall were above 91%, “Good” if both were above 80%, and flagged as “Notable Over- or Underestimation” when performance was imbalanced or mismatched with the ground truth. Only segmentations rated “Excellent” or “Good” were used in downstream analysis, helping ensure that all reported tissue patterns were based on high-confidence predictions.

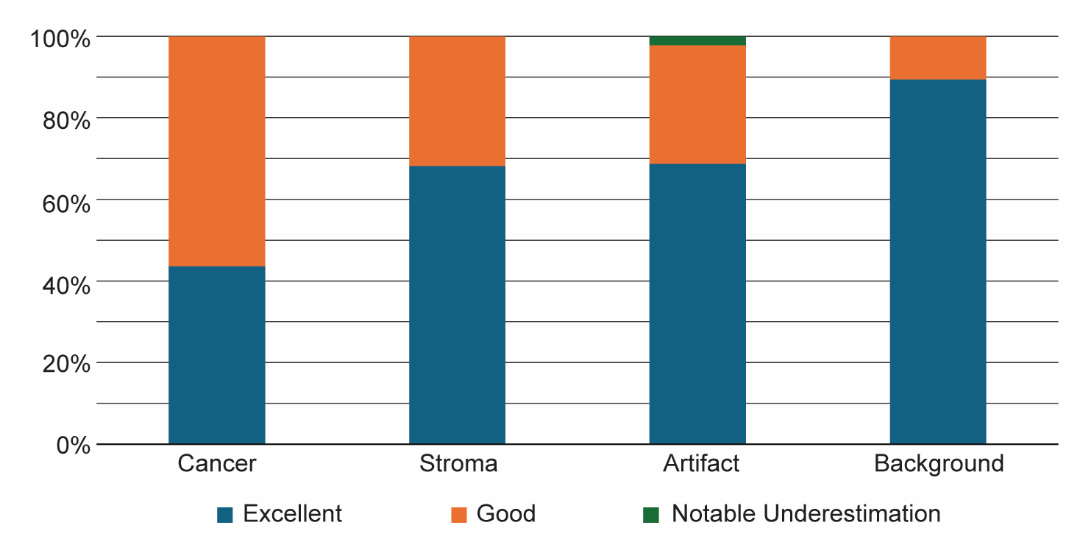
